# Rapid sequence evolution is associated with genetic incompatibilities in the plastid Clp complex

**DOI:** 10.1101/2021.07.13.452280

**Authors:** Salah E. Abdel-Ghany, Lisa M. LaManna, Zora Svab, Haleakala T. Harroun, Pal Maliga, Daniel B. Sloan

**Affiliations:** Department of Biology, Colorado State University, Fort Collins, CO 80523, USA; Waksman Institute of Microbiology, Rutgers University, Piscataway, NJ 08854, USA; Department of Plant Biology, Rutgers University, New Brunswick, NJ 08901, USA

**Keywords:** *clpP*, cytonuclear coevolution, epistasis, *Nicotiana*, plastome editing, *Silene*

## Abstract

The plastid caseinolytic protease (Clp) complex plays essential roles in maintaining protein homeostasis and comprises both plastid-encoded and nuclear-encoded subunits. Despite the Clp complex being retained across green plants with highly conserved protein sequences in most species, examples of extremely accelerated amino acid substitution rates have been identified in numerous angiosperms. The causes of these accelerations have been the subject of extensive speculation but still remain unclear. To distinguish among prevailing hypotheses and begin to understand the functional consequences of rapid sequence divergence in Clp subunits, we used plastome transformation to replace the native *clpP1* gene in tobacco (*Nicotiana tabacum*) with counterparts from another angiosperm genus (*Silene*) that exhibits a wide range in rates of Clp protein sequence evolution. We found that antibiotic-mediated selection could drive a transgenic *clpP1* replacement from a slowly evolving donor species (*S. latifolia*) to homoplasmy but that *clpP1* copies from *Silene* species with accelerated evolutionary rates remained heteroplasmic, meaning that they could not functionally replace the essential tobacco *clpP1* gene. These results suggest that observed cases of rapid Clp sequence evolution are a source of epistatic incompatibilities that must be ameliorated by coevolutionary responses between plastid and nuclear subunits.

## INTRODUCTION

Owing to the endosymbiotic origins of mitochondria and plastids (chloroplasts) and the subsequent history of cytonuclear integration in eukaryotes, many of the key enzyme complexes in these organelles are assembled from a mix of subunits encoded in both nuclear and cytoplasmic genomes (Rand, et al. 2004; Gould, et al. 2008; Gray 2012; Roger, et al. 2017; Sloan, et al. 2018). As such, core eukaryotic functions require coordinated regulation and coevolution between genomes that differ in nearly every respect, including their modes of transmission, mutation rates, copy numbers, and mechanisms of replication and expression.

The plastid caseinolytic protease (Clp) is one example of a cytonuclear enzyme complex. In the model angiosperm *Arabidopsis thaliana*, the proteolytic core of this complex consists of 14 subunits in two heptameric rings encoded by nine different genes, including eight nuclear loci and a single gene (*clpP1*) in the plastid genome (plastome). The complete Clp complex also contains a number of associated chaperones and adapters that are all nuclear-encoded (Nishimura, et al. 2015; Nishimura and van Wijk 2015). The *clpP1* gene appears to be essential based on studies that have modified the plastome to knockout this gene in the angiosperm *Nicotiana tabacum* (Shikanai, et al. 2001; Kuroda and Maliga 2003) and the green alga *Chlamydomonas reinhardtii* (Huang, et al. 1994). Likewise, disruption of nuclear genes that encode core Clp subunits also produces severe phenotypic effects (Sjögren, et al. 2006; Zheng, et al. 2006; Koussevitzky, et al. 2007; Kim, et al. 2009; Kim, et al. 2013; Moreno, et al. 2017). The Clp complex plays a central role in protein quality control and homeostasis in plastids, and many potential proteolytic targets have now been identified (Majeran, et al. 2000; Nishimura, et al. 2013; Tapken, et al. 2015; Apitz, et al. 2016; Pulido, et al. 2016; Moreno, et al. 2018; Welsch, et al. 2018; Wu, et al. 2018; Montandon, et al. 2019).

In accordance with the key functions of the Clp complex, the *clpP1* gene is almost universally retained in the plastomes of green plants and is highly conserved in sequence in most species (Williams, et al. 2019). For example, angiosperm ClpP1 protein sequences often share ∼60% amino acid identity with their counterparts in cyanobacteria despite more than a billion years of evolutionary divergence. However, a number of independent angiosperm lineages exhibit massive increases in rates of protein sequence divergence in this protein (Erixon and Oxelman 2008; Williams, et al. 2019). In extreme cases, ClpP1 sequences have been found to retain less than 35% amino acid identity even between closely related species in the same genus (Rockenbach, et al. 2016). The level of sequence divergence can be so extensive that the *clpP1* gene is often overlooked in plastome annotations (Haberle, et al. 2008; Straub, et al. 2011; Fajardo, et al. 2013; Yao, et al. 2015) or predicted to be a nonfunctional pseudogene (Hirao, et al. 2008; Zhang, et al. 2014). However, in the limited number of cases that have been investigated to date, these highly divergent *clpP1* gene copies appear to be functional, with evidence of transcription and proper splicing (Williams, et al. 2015) or even translation and likely assembly into the core Clp complex (Williams, et al. 2019). Although the relative contributions of mutation and selection to these changes in rates of *clpP1* evolution remain unclear, there is evidence that the gene may be subject to extensive positive selection for amino-acid substitutions in some species (Erixon and Oxelman 2008).

In angiosperms with highly divergent copies of *clpP1*, there are also correlated increases in the rate of protein sequence evolution in nuclear-encoded Clp subunits (Williams, et al. 2019; Forsythe, et al. 2021) and signatures of positive selection on these nuclear genes based on both population genetic and phylogenetic data (Rockenbach, et al. 2016). The apparent action of positive selection on interacting proteins encoded in two different genomes is reminiscent of antagonistic molecular coevolution that can occur between hosts and pathogens (Hughes and Nei 1988; Aguileta, et al. 2009). As such, it has been hypothesized that the examples of extreme protein sequence divergence in the Clp complex could be driven by selfish plastid-nuclear interactions (Rockenbach, et al. 2016). Although the origins of mitochondria and plastids represent some of the most intimate and important mutualisms in the history of life, they can still be involved in antagonistic interactions with the nucleus, including selfish over-replication within cells and conflict over allocation to female vs. male reproduction (Havird, et al. 2019). Although these types of conflicts have been more extensively documented in mitochondria, plastids may also be involved in selfish interactions with the nucleus. For example, the proliferation of genetically incompatible plastids within heteroplasmic *Oenothera* lines has been associated with variation in plastid-encoded components of fatty acid biosynthesis pathways and intracellular competition (Sobanski, et al. 2019). Plastid-nuclear incompatibilities have also been implicated in male sterility (Bogdanova, et al. 2015; Nováková, et al. 2019), which is a classic source of cytonuclear conflict (Touzet and Budar 2004; Fujii, et al. 2011).

Alternatively, positive selection could be indicative of some combination of adaptation, mutation accumulation, and compensatory coevolution rather than an antagonistic interaction. For example, in some systems, cytoplasmic genomes may accumulate disruptive sequence changes that require compensatory changes in nuclear genes to maintain function (Osada and Akashi 2012; Sloan, et al. 2017). Generation of “mismatched” combinations of plastid and nuclear genotypes through crossing designs or protoplast fusion have often identified plastid-nuclear incompatibilities (Greiner, et al. 2011; Barnard-Kubow, et al. 2016), and in rare cases those incompatibilities have been traced to specific loci (Schmitz-Linneweber, et al. 2005; Bogdanova, et al. 2015; Zupok, et al. 2020). A more targeted approach to probing plastid-nuclear incompatibilities involves genome editing to manipulate or replace individual genes. For example, plastome transformation in tobacco (*N. tabacum*) to replace its copy of *rbcL* with the orthologous sequence from sunflower (*Helianthus annuus*) successfully generated a hybrid tobacco-sunflower Rubisco complex, which contains plastid-encoded RbcL subunits and nuclear-encoded RbcS subunits (Kanevski, et al. 1999). Although catalytically active, this hybrid complex exhibited reduced function and evidence of incompatibilities between RbcL and RbcS subunits derived from different angiosperm lineages.

Here, we apply a plastome editing approach to replace tobacco *clpP1* with copies from other angiosperms with highly divergent rates of *clpP1* sequence evolution. For donor species, we take advantage of the extreme rate variation within the angiosperm genus *Silene* (Caryophyllaceae). Some species within this genus, including *S. latifolia*, have retained the typically low rates of ClpP1 protein sequence evolution found in most angiosperms, whereas others such as *S. conica* and *S. noctiflora* exhibit massive rate accelerations (Erixon and Oxelman 2008; Sloan, Triant, Forrester, et al. 2014; Rockenbach, et al. 2016). These *Silene* species are separated from tobacco by the same amount of divergence in time but differ radically in levels of ClpP1 protein sequence divergence (Figure 1). If the high levels of ClpP1 divergence in species such as *S. conica* and *S. noctiflora* reflect the role of this protein in some type of selfish plastid phenotype, introducing their gene sequences into tobacco would be akin to exposing a naïve host to a pathogen. Under this model, we would predict that these transgenic genotypes would exhibit signs of selfish conflict such as over-replication of the plastome within cells or induction of male sterility phenotypes. On the other hand, if the rapid sequence changes in Clp subunits solely reflect mutualistic cytonuclear coevolution to maintain compatibility, then increasing levels of sequence divergence should be associated with greater levels of epistatic incompatibilities and loss of Clp function when transformed into tobacco.

**Figure 1.**
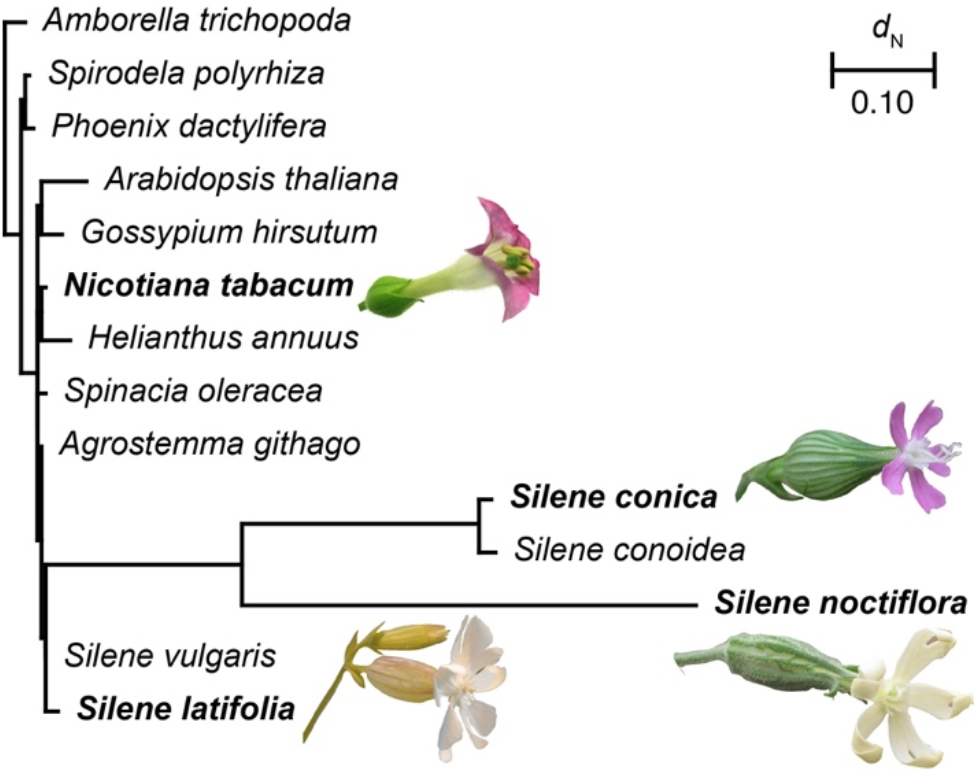
Variation in *clpP1* nonsynonymous substitutions rate (*d*_N_) in *Silene* species relative to a sample of diverse angiosperms with relatively conserved copies of *clpP1*. Branch lengths were estimated with codeml within the PAML v4.9j package (Yang 2007), using a constrained topology. *Silene noctiflora* was constrained to be sister to the *S. conica*/*conoidea* lineage, although support for this relationship has been weak or inconsistent in previous studies (Rautenberg, et al. 2012; Havird, et al. 2017; Jafari, et al. 2020). The species used in this study are highlighted in bold text and accompanied by a flower image.

## MATERIALS AND METHODS

### Construction of plastid transformation vectors for replacement of *clpP1* in tobacco

To replace the native *clpP1* gene in the tobacco plastome with orthologous sequences from donor species, we relied on particle bombardment of leaf tissue and the natural homologous recombination activity within plastids (Svab and Maliga 1993). This strategy involved combining the desired donor *clpP1* sequence with a selectable marker and flanking sequences from tobacco plastome that would serve as substrate for homologous recombination (Figure 2). A region of the plastome containing 958 bp of the *psbB* gene and intergenic region between *clpP1* and *psbB* (corresponding to plastome positions 74,728-75,686; GenBank accession KU199713.1) was amplified by PCR as a *BamHI*/*SmaI* fragment using psbBHR-S and psbBHR-AS primers (Table S1). The PCR product was cloned in a *BamHI/SmaI*-linearized pBluescript KS (+) phagemid (Stratagene) producing pBS-*v*1 (Figure 2). Also, a 3200-bp plastome fragment containing the *clpP1* operon and intergenic region of *clpP1* and *psbB* (corresponding to plastome position 71,528-74,728) was amplified as a *XhoI*/*SmaI* fragment using ClpP1HR-S and ClpP1HR-AS primers (Table S1) and cloned in pBS-*v*1 to produce pBS-*v*2. The chimeric spectinomycin resistance (aa*dA*) construct with 16S rRNA promoter (Prrn) and *psbA* 3’ regulatory regions (Svab and Maliga 1993) was synthesized and cloned into the *SmaI* site in pBS-*v*2 to produce pBS-*v*3. The orientation of the insert was selected such that the transcription of *aadA* is in the opposite direction of the *clpP1* operon. A *SphI* restriction site was introduced behind *SmaI* at the 5’ end of the *aadA* construct for exchange of the native tobacco *clpP1* with counterparts from donor species using a unique *BstZ17I* restriction site in the *clpP1* operon. Coding sequences from *clpP1* cDNA from donor species (tobacco, *S. conica, S. latifolia*, and *S. noctiflora*) were commercially synthesized (GenScript, Piscataway, NJ) with the native tobacco regulatory elements and an introduced *SphI* restriction site and then cloned into pBS-*v*3, yielding pBS-*v*4, pBS-*v*6, pBS-*v*7 and pBS-*v*9, respectively), using *SphI* and *BstZ17I* cloning sites (Figure 2). Note that most angiosperms have two introns in *clpP1*, so the absence of these introns in the cDNA donors represented a difference relative to the genomic sequence (although the introns have already been lost from the plastomes of *S. conica* and *S. noctiflora*). In addition, *S. latifolia* has a C-to-U RNA editing site in codon 187 that is widespread in angiosperms but has been lost in *S. conica, S. noctiflora*, and *Nicotiana* (Williams, et al. 2019). Therefore, for the *S. latifolia* cDNA construct, we used a T at this position to mimic the edited state. Plasmids were transformed into DH5α *E. coli*, propagated, extracted by QIAGEN Plasmid Midi kit, and used for plastome transformation.

**Figure 2.**
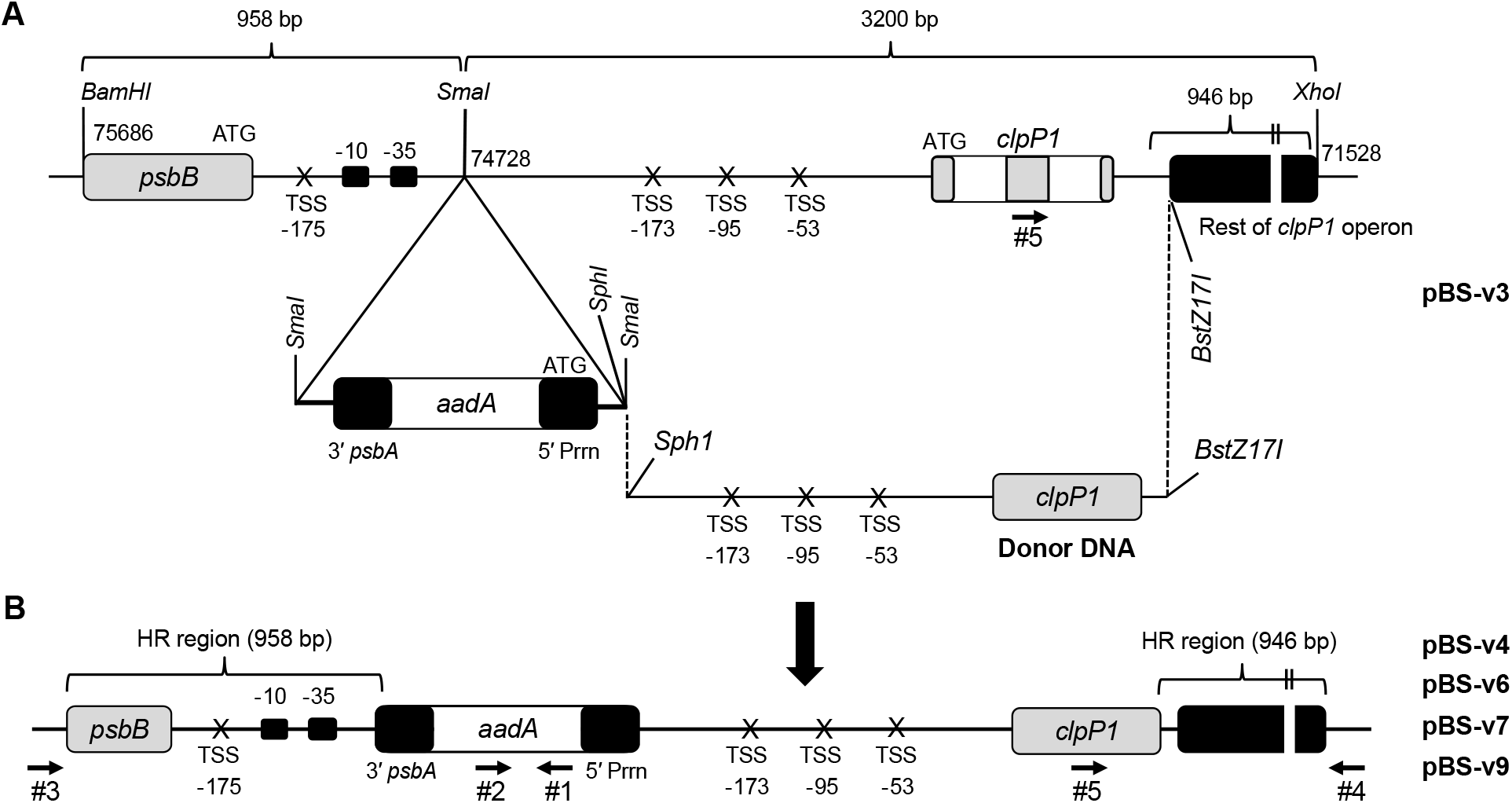
Targeted replacement of *clpP1* in the tobacco plastome with cDNA counterparts from different *Silene* species. (A) Physical map of the targeting region in the plastome of wild type tobacco cloned in pBluescript KS (+) plasmid (pBS-v3, see text for details). The chimeric selectable marker gene *aadA* conferring resistance to spectinomycin and streptomycin is driven by the rRNA operon promoter (5’ *Prrn*). The 3’ untranslated region of the *psbA* gene (3’ *psbA*) was added to stabilize the mRNA (Svab and Maliga 1993). (B) Plastid-targeting region with flanking homologous recombination sequences (HR regions), donor *clpP1* coding sequences, and *aadA* cassette. The *aadA* cassette was introduced in antisense direction relative to *clpP1* operon. Restriction sites used for cloning and replacement are indicated. Numbered horizontal arrows represent primers used in PCR validation of insertion and orientation (Table S1). Transcription start sites (TSSs) are indicated in the intergenic regions between the *psbB* gene and *clpP1* operon (Hajdukiewicz, et al. 1997). Certain regions of the map are expanded and not drawn to scale for the sake of clarity.

### Plastid transformation and selection of transplastomic tobacco lines

Tobacco (*N. tabacum* cv. Petit Havana) plants were grown in a controlled growth chamber at 250 μmol photons m^-2^ s^-1^ light intensity (16 h day, 25-27 °C). Plastid transformation was performed according to (Svab and Maliga 1993), using the biolistic PDS-1000/He particle bombardment system (Bio-Rad). Young leaves of aseptically grown tobacco plants were bombarded with plasmid-DNA-coated 0.6 μm gold particles (Bio-Rad) and selected on RMOP media with 500 μg ml^-1^ spectinomycin (Svab and Maliga 1993). The primary spectinomycin-resistant shoots were confirmed by testing for double resistance on RMOP medium containing spectinomycin and streptomycin (500 μg ml^-1^ each) (Svab and Maliga 1993; Bock 2001). Transplastomic plants were subjected to 4 or 5 additional rounds of regeneration (Table S2) in the presence of spectinomycin to enrich the transgenic plastome and to select against the wild type genome copies. Correct transgene insertion and orientation were confirmed by PCR, and relative abundance of wild type and modified plastomes was assessed by quantitative PCR (qPCR) and DNA gel (Southern) blotting (see below).

Following the antibiotic-selection regime described above, we removed the transplastomic lines from selection to assess the stable maintenance of the edited plastomes. Using at least three biological replicates for each of the *Silene* or tobacco control *clpP1* replacements, we maintained parallel cultures both with and without spectinomycin through two additional rounds of subculture and regeneration.

### PCR and qPCR assays of modified plastomes

Total cellular DNA was prepared from regenerated leaves by the method of Doyle and Doyle (1990) and quantified by Qubit 2.0 Fluorometer (Invitrogen), using the dsDNA HS kit. PCR amplification was carried out using EmeraldAmp GT PCR Master Mix (Clontech, Takara) according to the manufacturer’s protocol and the primers shown in Table S1 and Figure 2. PCR conditions were 2 min at 98 °C followed by 35 cycles of 98 °C for 10 s, 60 °C for 30 s and 72 °C for a time period that depended on the length of the target PCR product. PCR amplicons were electrophoresed on 1% (w/v) agarose gels. For testing the heteroplasmic levels of wild type and transgenic *clpP1* copies by qPCR, 0.1 ng total cellular DNA and 0.25 μM sense and antisense primers were used in a 10 μl reaction with iTaq Universal SYBR Green Supermix (Bio-Rad). The reaction volumes were heated at 95 °C for 3 min, followed by 40 cycles of 95 °C for 10 s and 60 °C for 30 s and a subsequent melt curve analysis on a Bio-Rad CFX96 Touch thermal cycler and real-time PCR detection system. QClpP1-F and QClpP1-R primers (Supplementary Table 1) located in exon 2 and intron 1, respectively were used in qPCR to estimate the copy number of wild type tobacco genomic *clpP1* (hereafter referred to as the exon/intron marker). The *aadA* transgene and the single-copy *psbA* gene were used as references for normalization. The amplification efficiency was calculated from the slope of a standard curve obtained from 2-fold serial dilution of genomic DNA ranging from 5 ng to 0.078 ng.

For each of the four cDNA constructs (tobacco, *S. conica, S. latifolia*, and *S. noctiflora*), six biological replicates were analyzed for each qPCR marker. Where possible, we chose independent transformants for these replicates. However, in some cases there were fewer than six independent transformant lines with the correct insert (five for *N. tabacum*, four for *S. conica*, and three for *S. latifolia*). In these cases, we included two subcultures from the same primary transformant to obtain the desired number of biological replicates. Each reaction was run in duplicate (technical replicates), and the threshold cycle (C_t_) values for these two replicates were averaged for further analysis. After removing lines from selection (see above), these qPCR assays were then repeated on parallel cultures growing with and without spectinomycin.

We used a ΔC_t_ approach to estimate the relative abundance of wild type vs. transgenic plastome copies by comparing the C_t_ value for the exon/intron marker to the C_t_ values of each of the two aforementioned reference markers (*aadA* and *psbA*). Because the exon/intron maker should only amplify in wild type copies and the *aadA* markers should only amplify in transgenic copies, a ΔC_t_ value (i.e., C_t [exon/intron]_ – C_t [*aadA*]_) of 0 would indicate 50/50 heteroplasmy, whereas positive ΔC_t_ values for this comparison would indicate an excess of transgenic copies and negative values would indicate an excess of wild type copies. The interpretation of comparisons to the *psbA* reference marker (i.e., ΔC_t_ = C_t [exon/intron]_ – C_t [*psbA*]_) is similar except that the *psbA* marker should amplify in both wild type and transgenic plastomes. Therefore, for this comparison the 50/50 heteroplasmy point should correspond to a C_t_ value of 1 rather than 0. To test for significant difference among the four transgenic lines, a one-way ANOVA was performed with the aov function in R v3.6.3, followed by *post hoc* pairwise comparisons with the TukeyHSD function.

### Southern blotting

Total genomic DNA extracted from wild type and regenerated leaves of transplastomic lines was digested with *NruI* and *NarI* restriction enzymes, electrophoresed in a 0.8% (w/v) agarose gel, transferred to a nylon membrane by capillary transfer, and UV cross-linked to the membrane (Stratagene). Restriction enzymes were selected to produce two fragments (5211 and 2132 bp) from the wild type plastome and one fragment (∼7 kb) from transplastomes (with slight differences in length depending on the length of the donor *clpP1* transgene), as the *NarI* restriction site is located inside the second intron of *clpP1* genomic sequence but not present in any of the cDNA transgenes. Two probes (probe 1 and 2) from wild type tobacco plastome sequence flanking *NarI* restriction site were amplified by PCR using primers described in Table S1 and labeled with Biotin DecaLabel DNA Labeling Kit (Thermo Scientific). Therefore, while wild type samples are expected to show two bands (∼2 and ∼5 kb), only one band (∼7 kb) is expected in homoplasmic transplastomic lines, and heteroplasmic lines should have all three bands (∼7, ∼5, and ∼2 kb). Membranes containing digested DNA from transplastomic plants and wild type tobacco were hybridized with denatured probes for 18 h at 42 °C, washed, and incubated with a streptavidin HRP-conjugated antibody following the protocol in the Thermo Scientific Pierce Chemiluminescent Nucleic Acid Detection kit.

### Materials Availability

The constructs used for plastome transformation have been deposited to Addgene are available under accessions 173794-173797.

## RESULTS AND DISCUSSION

### Editing of tobacco plastome to replace native *clpP1* sequence with *Silene* counterparts

Using antibiotic selection, we confirmed that biolistic delivery was successful in introducing the *aadA* marker into the tobacco plastome as part of each of the four *clpP1* donor constructs (Figure 2). Growth on spectinomycin plates identified numerous potential transformants. Because spectinomycin resistance can arise by spontaneous mutations in plastid rRNA sequence (Svab and Maliga 1993), dual selection with spectinomycin and streptomycin was conducted to distinguish spontaneous mutants from lines with true transgene-mediated resistance (Svab, et al. 1990). On selection media, the calli of the resistant clones are green, while the sensitive ones are white (Maliga, et al. 1990).

PCR screening with multiple primers pairs (Figure 2 and Table S1) was used to confirm that double-resistant lines contained the full transgenic construct inserted in the expected location. The presence of the full-length transgene was also confirmed by Southern blot analysis for select lines (Figure S1). In some cases, lines carried the *aadA* selectable marker but exhibited evidence of rearrangements or internal recombination events (examples shown Figure S2), which is not surprising given the highly recombinational nature of plastomes. We restricted our analysis to lines that produced PCR products with expected lengths for markers spanning the boundary between the insert and flanking sequences on each side (Figure 3). Using this approach, we identified lines derived from at least three independent transformation events for each of the four *clpP1* donor constructs (tobacco, *S. conica, S. latifolia*, and *S. noctiflora* cDNAs). Where necessary, we propagated lines such that we had at least six biological replicates for each construct (Table S2).

**Figure 3.**
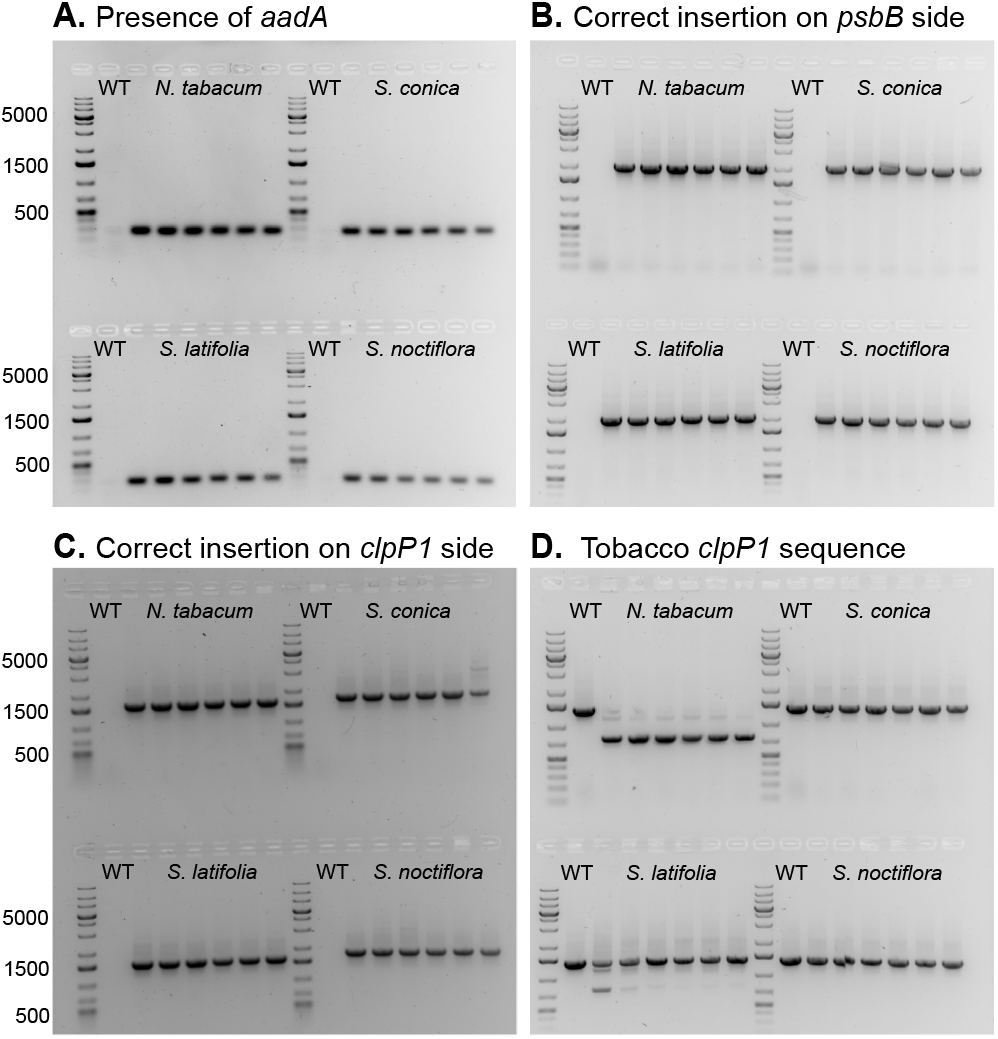
Confirmation of proper insertion and orientation of *clpP1* replacement constructs. Each gel shows PCR products from amplification of total-cellular DNA extracted from wild type (WT) tobacco and the six replicate transgenic lines for each of the four *clpP1* cDNA replacement constructs (indicated by the name of the donor species; Table S2). (A) Detection of *aadA* selectable marker gene with primers #1 and #2; expected product size 276 bp. (B) Confirmation of proper insertion of *aadA* transgene with flanking sequence on the *psbB* side, using primers #1 and #3; expected product size 2002 bp. (C) Confirmation of proper insertion of *aadA* transgene with flanking sequence on the *clpP1* side, using primers #2 and #4; expected product size 1554 bp. (D) Amplification of *clpP1* and flanking sequence with a tobacco-specific primer (#5) and primer #4; expected product size of 1400 bp for native genomic DNA and 776 bp for cDNA. The *Silene* cDNA constructs are not expected to amplify with this marker except for *S. latifolia*, which appears to show weak cross-reactivity presumably because of its greater degree of sequence similarity with tobacco. Sizes of the dark marker bands (in bp) are shown to the left. All primer numbers refer to Figure 2 and Table S1.

### More divergent *clpP1* donor sequences were restricted to lower heteroplasmic frequencies

Typically, homoplasmic shoots are obtained after 3-5 months during the second or third cycle of regeneration in the presence of antibiotic (Svab and Maliga 1993). However, even after five cycles of regeneration, the frequency of wild type plastome copies detectable by qPCR was higher in the lines carrying *Silene clpP1* replacement constructs than in the control lines carry tobacco cDNA replacements, indicating that these *Silene* replacement lines had generally not reached homoplasmy (Figure 4). The measured primer efficiencies for all qPCR markers were very close to 100% (exon/intron: 102.0%; *aadA*: 97.9%; *psbA*: 98.0%). Therefore, ΔC_t_ comparisons should provide an accurate estimate of the relative abundance of modified vs. wild type plastome copies. Using tobacco *clpP1* cDNA itself to replace the corresponding tobacco genomic sequence was most successful, but even for this control construct, there was a wide range among lines for the estimated heteroplasmy levels (Table S3). The ΔC_t_ values for the tobacco cDNA lines imply that biological replicates ranged anywhere from a ∼4-fold to ∼30-fold excess of modified plastomes compared to wild-type, with the mean ΔC_t_ values indicating a ∼10-fold excess. The three *Silene* donor sequences all exhibited significantly lower levels of transgene enrichment than the tobacco cDNA control (Figure 4). However, there was a clear distinction between the more conserved *clpP1* sequence of *S. latifolia* and its highly divergent congeners, as the former obtained higher levels of transgene enrichment than either *S. conica* or *S. noctiflora*. The mean ΔC_t_ values for *S. latifolia* implied a ∼3.5-fold excess of the modified plastomes, whereas the *S. conica* and *S. noctiflora* lines both only appeared to reach roughly 50/50 heteroplasmy levels on average (Figure 4).

**Figure 4.**
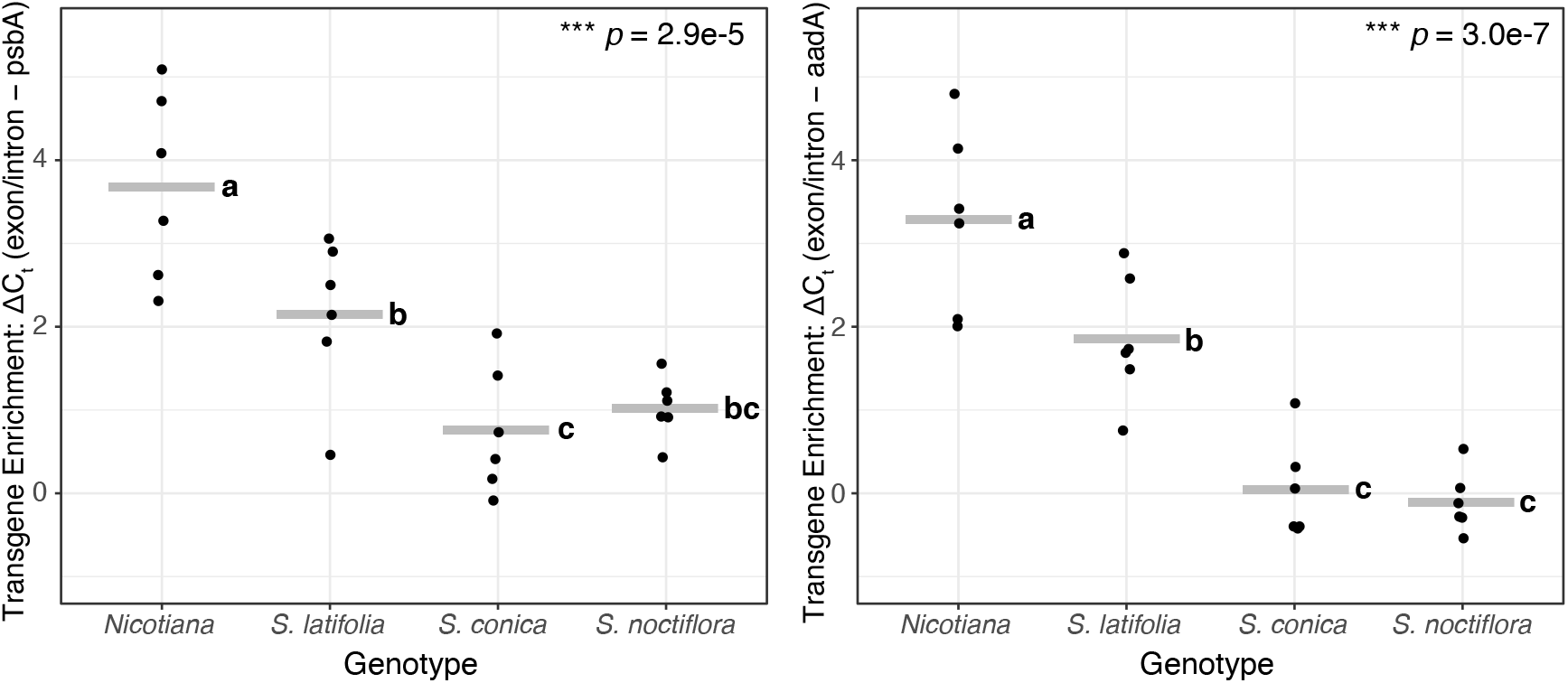
Varying levels of heteroplasmy obtained for different replacements of the native *clpP1* sequence in tobacco. Levels of enrichment by antibiotic selection for the transgenic plastomes were inferred by qPCR analysis, comparing a wild type marker (spanning an exon/intron junction in the native *clpP1*) against a shared reference gene (*psbA*; left) or the antibiotic resistance maker in the transgenic cassette (*aadA*; right). Higher ΔC_t_ values are indicative of reduced abundance of the wild type maker and greater enrichment of transgenic plastomes. Each black point represents a biological replicate (averaged from two technical replicates), and gray bars indicate the mean value for that donor sequence. Statistical significance was assessed with a one-way ANOVA. Lowercase letters indicate significant pairwise differences as identified by *post hoc* Tukey’s HSD tests.

Like most plants, tobacco is known to harbor numerous insertions of plastid DNA in its nuclear genome, which are known as “nupts” (Rousseau-Gueutin, et al. 2011). Therefore, it is likely that some of the wild type signal detected by qPCR actually results from amplifying nupts rather than true plastome copies. Unfortunately, a complete accounting of nupts is not available for *N. tabacum*, as insertions of organellar DNA can be very difficult to accurately identify even for some of the highest quality nuclear genome assemblies (Stupar, et al. 2001). However, because plastomes typically occur in hundreds to thousands of copies per cell (Greiner, et al. 2020), the signal from nupts should be relatively weak compared to true plastid DNA. Moreover, the same nuclear background was used for all lines, so any contribution from nupts should be roughly equal across all samples. Nevertheless, the presence of nupts can make for an ambiguous distinction between homoplasmy and heteroplasmy in qPCR data. Therefore, we subjected transgenic lines to further subculturing and took additional steps to test for homoplasmy as described below.

### Homoplasmic replacement of native *clpP1* obtained for tobacco cDNA control and *S. latifolia clpP1* but not for *S. noctiflora* and *S. conica* donors

Using the transgenic lines described above, we established parallel cultures both with and without antibiotic selection and assayed the relative abundance of transgene (*aadA*) and wild-type (exon/intron) markers by qPCR (Figure 5, Table S4). Lines harboring tobacco and *S. latifolia* cDNA constructs maintained high transgene frequencies both with and without selection (Figure 5), raising the possibility that they had reached homoplasmy. In the absence of antibiotic selection, we would expect transgenes that were still heteroplasmic to reduce in frequency, especially if *aadA* expression is costly or the *clpP1* replacement is not fully functional in a tobacco genetic background. In contrast to the tobacco and *S. latifolia* cDNA constructs, the lines containing *S. noctiflora* and *S. conica* donor sequences exhibited much lower transgene frequencies, indicative of heteroplasmy. As expected, the *S. noctiflora* lines generally showed a further reduction in transgene frequency when removed from antibiotic selection. Surprisingly, this was not the case for the *S. conica* lines, which remained at approximately the same frequencies regardless of whether they were cultured with or without antibiotics (Figure 5).

**Figure 5.**
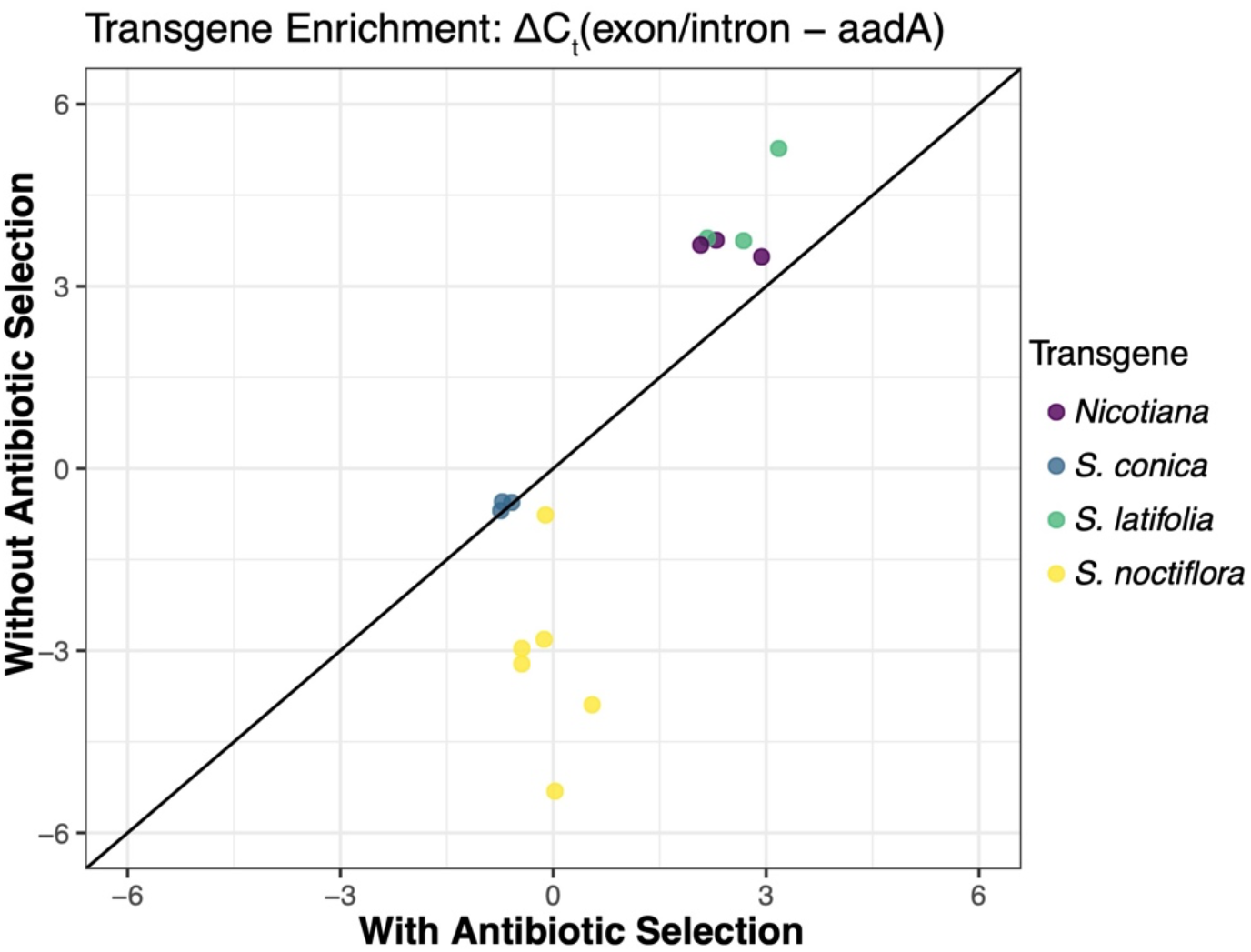
qPCR analysis of transplastomic lines grown in parallel cultures either with or without antibiotic selection. Each point represents a transplastomic line and the corresponding transgene enrichment values under the two selection regimes. qPCR analysis compared a wild type marker (spanning an exon/intron junction in the native *clpP1*) against the antibiotic resistance maker in the transgenic cassette (*aadA*). Higher ΔC_t_ values are indicative of reduced abundance of the wild type maker and greater enrichment of transgenic plastomes. The black diagonal line represents the 1:1 line. Points falling below that line indicate that the transgene dropped in frequency in the culture propagated without antibiotic selection.

A Southern blot analysis helped explain this surprising qPCR result for the *S. conica* lines (Figure 6). Although they were originally confirmed to contain the full transgenic construct (Figure 3), we found that these *S. conica* lines had since undergone a recombination event between wild type and transgenic plastomes. Therefore, rather than being heteroplasmic, they were homoplasmic for a recombinant plastome that appeared to contain both the *aadA* selection marker and the wild type genomic *clpP1* sequence (resulting in the false appearance of a ∼50/50 heteroplasmy in qPCR data). In contrast, the *S. noctiflora* lines still exhibited heteroplasmy for the wild type and transgenic plastomes, but most of these lines also showed evidence of recombinant haplotypes as well (Figure 6). Therefore, it appears that persistent heteroplasmy resulting from transgenic *clpP1* donor sequences that cannot replace the functionality of the native *clpP1* gene provides an ongoing opportunity for recombination between wild type and transgenic plastomes that can resolve the conflicting selection pressures for *clpP1* function and antibiotic resistance.

**Figure 6.**
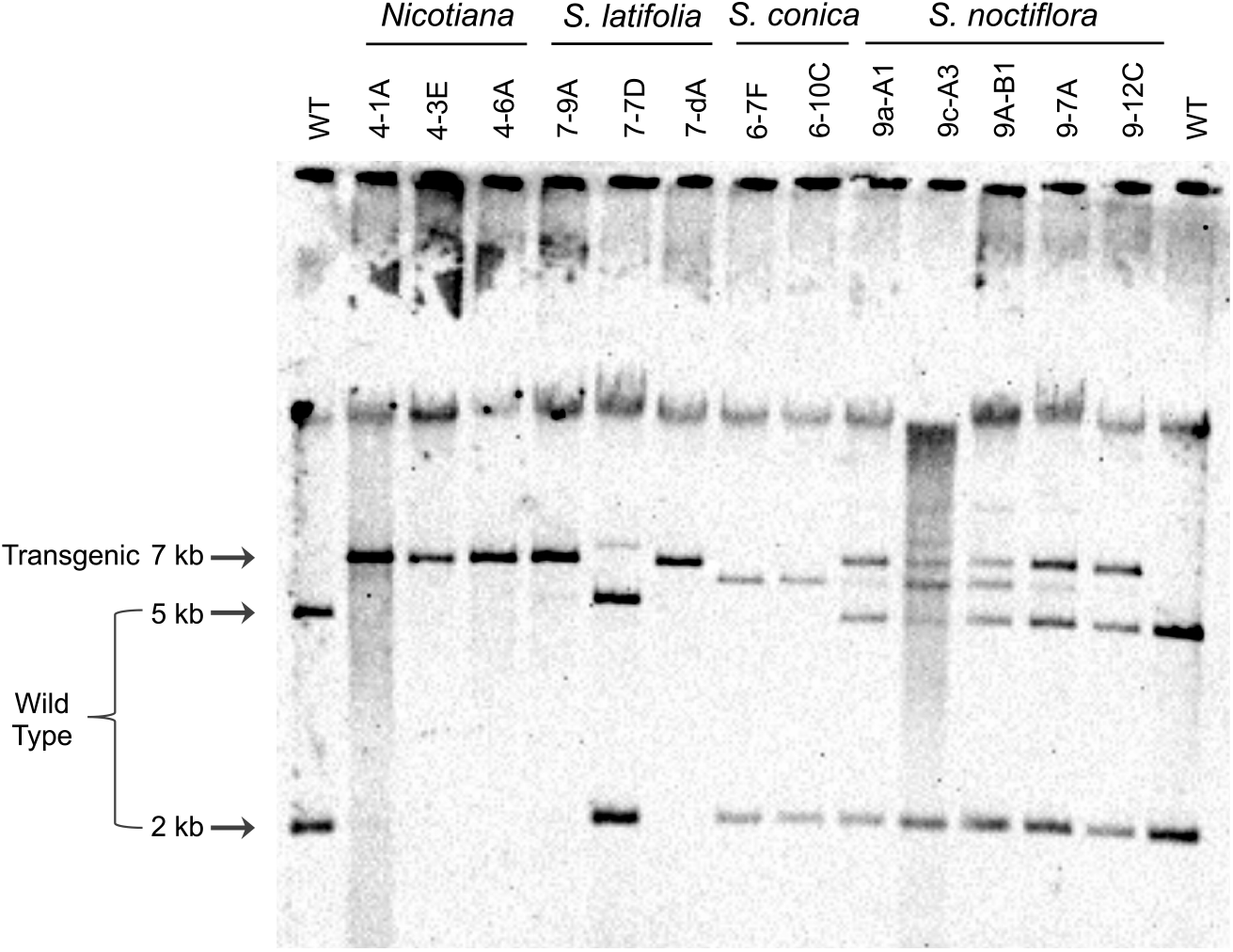
Southern blot analysis in which transplastomes should produce a single 7-kb band while wild type plastomes should produce both 5-kb and 2-kb bands as indicated. All three tobacco cDNA lines and the *S. latifolia* 7-dA line appear to be homoplasmic for the transplastome based on the absence of other detectable bands. All other lines are heteroplasmic for wild type and transgenic plastome, contain a recombinant haplotype, or both. The analysis was performed on samples maintained on antibiotic selection after parallel cultures were set up both with and without antibiotics (see main text).

The Southern blot analysis supported the inference that all three analyzed lines carrying the tobacco cDNA transgene were homoplasmic (Figure 6). In contrast, only one of the three analyzed *S. latifolia* lines (7-dA) appeared homoplasmic for the full transgenic construct in the Southern blot analysis. We confirmed with Sanger sequencing that this line retained the full-length *S. latifolia clpP1* coding sequence. Another *S. latifolia* line (7-7D) appeared homoplasmic for a haplotype with an internal recombination event, whereas a third line (7-9A) appeared nearly homoplasmic for the full transgene but also contained a faintly detectable recombinant haplotype. The fact that only one of the analyzed *S. latifolia* lines appears to have reached homoplasmy is consistent with the qPCR results showing that the tobacco cDNA transgenes are generally easier to drive to high frequency than the foreign *S. latifolia* transgene (Figure 4).

### Rapid *clpP1* evolution and the generation of epistatic plastid-nuclear incompatibilities

Our results support a model in which plastid-nuclear coevolution results in “matched” genotypes that are sensitive to disruption when novel genetic combinations are generated. Under the same selection conditions, *Silene*-derived *clpP1* transgenes were limited to lower heteroplasmic levels than the tobacco cDNA control, and this gap was more pronounced for *Silene* donors with histories of accelerated *clpP1* sequence evolution (Figure 4). Accordingly, additional antibiotic selection failed to drive the divergent *S. conica* and *S. noctiflora* transgenes to homoplasmy, but we were able to generate homoplasmic replacements for multiple tobacco cDNA control lines and a single *S. latifolia* line (i.e., the slowly evolving *Silene* donor; Figure 6). Previous analysis of chloroplast stromal protein fractions for *Silene* species with native gels and mass spectrometry has confirmed that the ClpP1 protein is still translated and appears to assemble with other subunits in the Clp proteolytic core (Williams, et al. 2019). In addition, the rates of ClpP1 evolution strongly correlate with amino-acid sequence divergence in interacting nuclear-encoded Clp subunits within *Silene* and more generally across angiosperms (Rockenbach, et al. 2016; Williams, et al. 2019). Taken together, these observations suggest that the lower relative abundance of the *Silene clpP1* transgenes (and especially those from *S. conica* and *S. noctiflora*; Figure 4) is caused by incompatibilities with the nuclear-encoded Clp subunits in tobacco. The apparent emergence of incompatibilities based on *clpP1* divergence among such close relatives is striking, especially given the observation that human (mitochondrial-targeted) *clpP* can partially substitute for the loss of its bacterial counterpart in *Bacillus subtilis* (Dittmar, et al. 2020) despite billions of years of evolutionary divergence. In that case, however, it should be noted that the Clp proteolytic core is homomeric (14 copies of the same subunit), in contrast to the highly heteromeric core of plastid Clp (Nishimura and van Wijk 2015).

Numerous studies have identified correlated rates of sequence evolution between genes in cytoplasmic and nuclear genomes and interpreted those as evidence of coevolution to maintain functional interactions (Osada and Akashi 2012; Sloan, Triant, Wu, et al. 2014; Pett and Lavrov 2015; Zhang, et al. 2015; Adrion, et al. 2016; Rockenbach, et al. 2016; Weng, et al. 2016; Zhang, et al. 2016; Havird, et al. 2017; Yan, et al. 2019; Forsythe, et al. 2021). Rarely, however, have such phylogenetic studies been coupled with functional tests for genetic incompatibilities. More generally, despite extensive evidence that hybridization can cause plastid-nuclear incompatibilities (Greiner, et al. 2011), the specific loci involved in those incompatibilities remain unclear except for a small number of cases (Schmitz-Linneweber, et al. 2005; Sobanski, et al. 2019; Zupok, et al. 2020). Our approach used here and previously (Kanevski, et al. 1999) to generate hybrid plastid-nuclear enzyme complexes with subunits derived from different species should be effective for testing for specific genetic incompatibilities.

We previously hypothesized that the rapid evolution and evidence for positive selection in *Silene* Clp complexes could reflect selfish dynamics and antagonistic coevolution between the plastome and nuclear genome (Rockenbach, et al. 2016). Under this hypothesis, we would predict that the divergent *clpP1* genes are acting as selfish elements that increase their own transmission at the expense of organismal fitness. Potential mechanisms could include intracellular advantages, such as preferential rates of replication/division of plastomes or plastids carrying certain *clpP1* variants, or organismal phenotypes such as male sterility that can boost transmission of maternally inherited organelle genomes (Havird, et al. 2019). We did not find any evidence for such a selfish advantage given the *S. conica* and *S. noctiflora clpP1* transgenes were unable to reach homoplasmy and were limited to lower heteroplasmic levels than other constructs. These findings do not rule out the possibility that some form of selfish dynamics contributed to accelerated Clp evolution in these lineages. However, they suggest that if any selfish benefit did exist, it is greatly outweighed by incompatibilities that accumulated over the cumulative history of sequence divergence and coevolution that is disrupted by putting these genes in a tobacco background. One avenue for future investigation would be to compare the levels of heteroplasmy obtained for the *S. conica* and *S. noctiflora* transgenes to simple transgenic knockouts of *clpP1*. Because *clpP1* is essential, knockouts do not reach homoplasmy in tobacco (Shikanai, et al. 2001; Kuroda and Maliga 2003). However, if they were able to reach higher heteroplasmic frequencies than the *Silene* replacements, it would indicate that the sequence divergence in *Silene* does not only lead to loss of function in a tobacco background but is actively harmful through dominant negative effects. In addition, the ability to introduce targeted modifications through site-directed mutagenesis creates the opportunity to dissect the specific changes within *clpP1* that are responsible for the observed incompatibilities. Such techniques are promising for unraveling the history of cytonuclear coevolution at a detailed molecular scale.

## ACKNOWLEDGEMENTS

We thank Alissa Williams for comments on an earlier version of this manuscript and Amber Torres and Matheus Fernandes Gyorfy for lab assistance. This work was supported by a grant from the National Science Foundation (MCB-1733227).

---

**Table S1.**
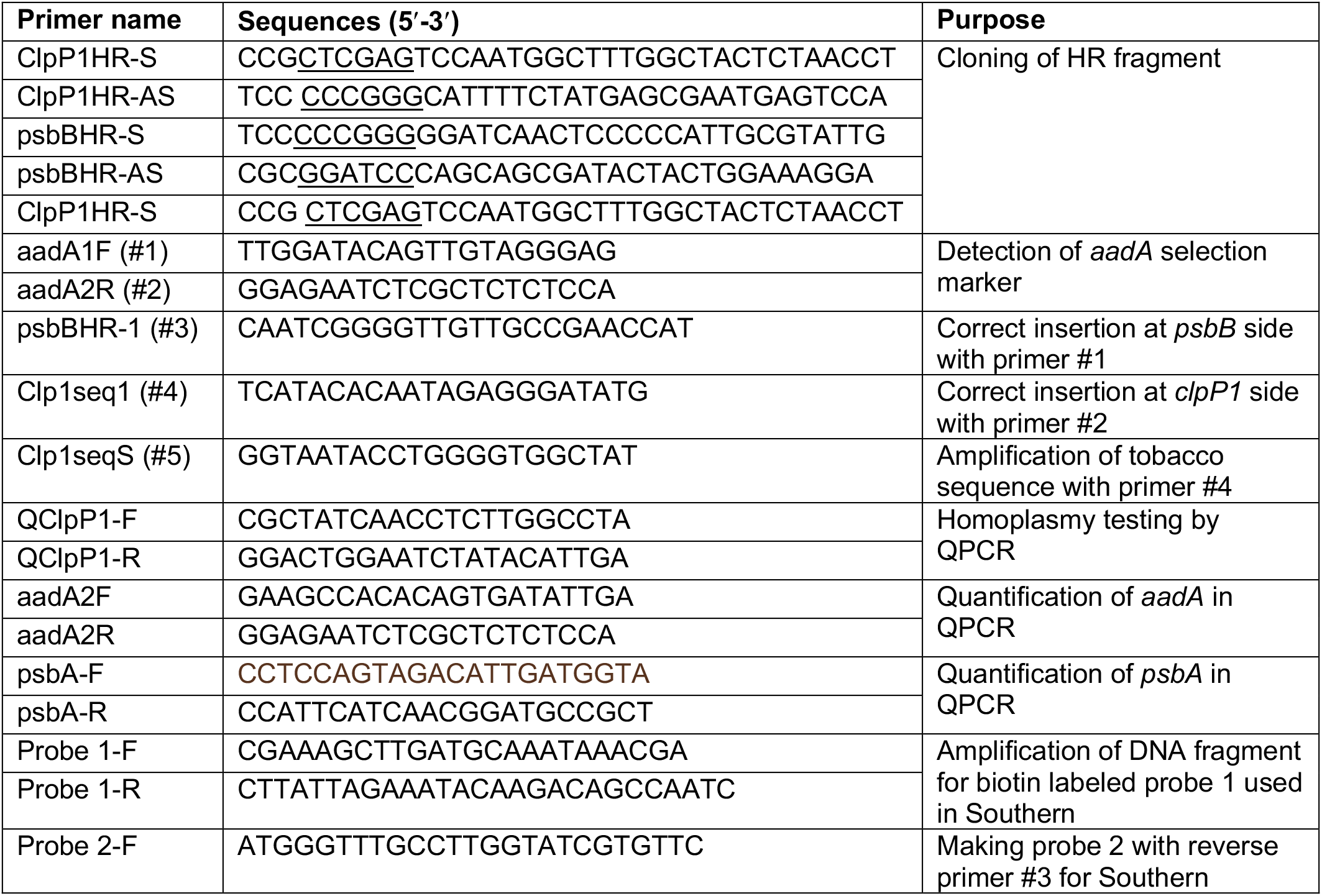
Primer sequences. Underlined sequences represent introduced restriction sites for cloning. Primer names parenthetically labeled #1 to #7 refer to locations described in Figure 2.

**Table S2.** Six biological replicates from each transplastomic cDNA replacement line. Pairs shown in cells without divider lines are subcultures derived from the same primary transformant. Values in parenthesis indicate the number of regeneration cycles of the different transplastomic lines in selection media.

| ***N. tabacum*** | ***S. conica*** | ***S. latifolia*** | ***S. noctiflora*** |
| --- | --- | --- | --- |
| 4-1A (5) | 6-3A (5) | 7-1B (5) | 9a-A (5) |
| 4-1B (5) | 6-3B (5) | 7-1C (5) | 9c-A (5) |
| 4-3E (5) | 6-5C (5) | 7-9A (5) | 9d-B (5) |
| 4-6A (5) | 6-7F (5) | 7-9B (5) | 9A-B (4) |
| 4-7B (5) | 6-10B (4) | 7d-A (4) | 9-7A (5) |
| 4b-A1 (5) | 6-10C (4) | 7d-C (4) | 9-12C (4) |
| 5 independent transformants | 4 independent transformants | 3 independent transformants | 6 independent transformants |

**Table S3.**
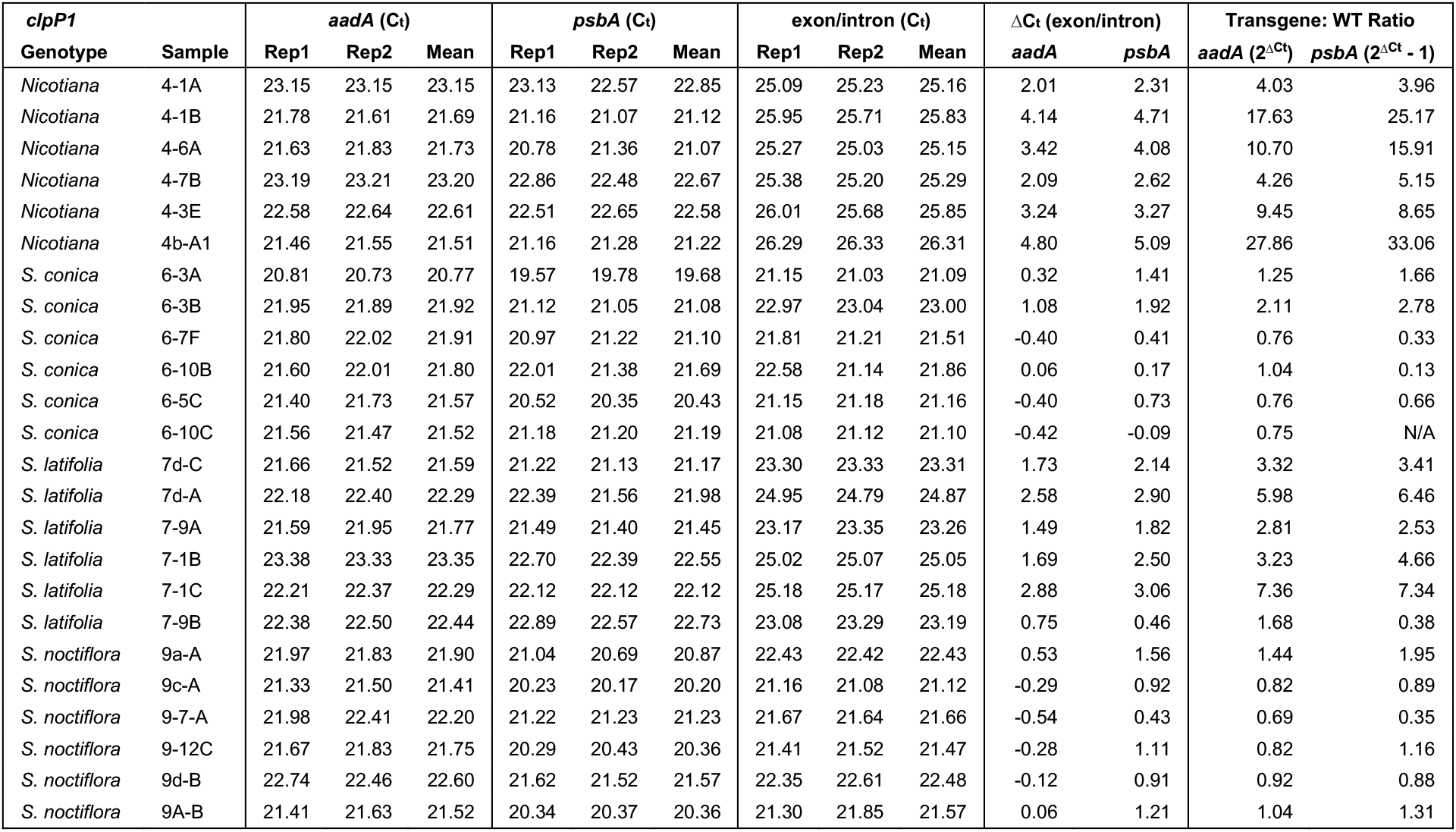
qPCR data from Figure 4 for estimating transgene enrichment relative to wild type plastomes

**Table S4.**
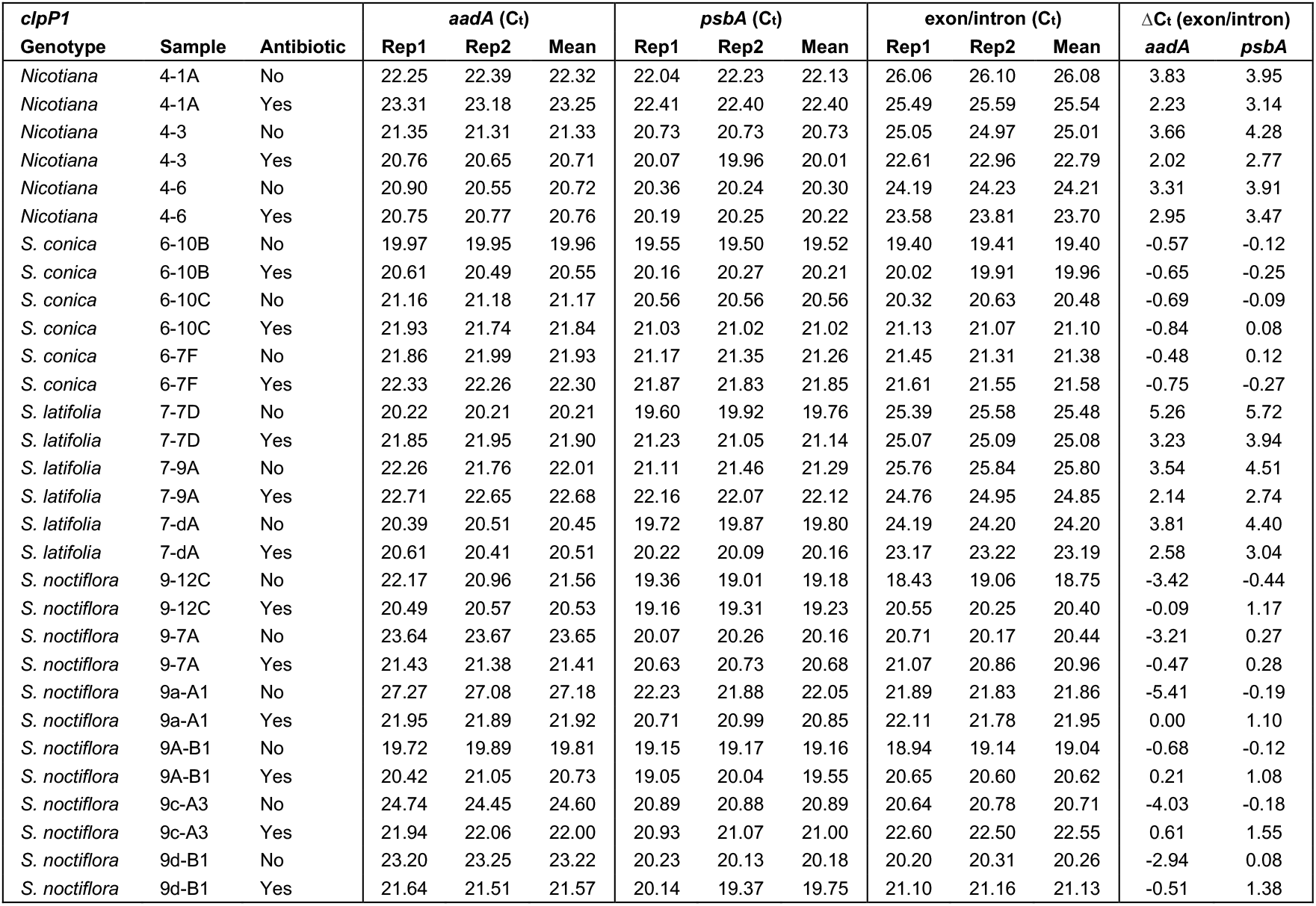
qPCR data from Figure 5 for estimating transgene enrichment relative to wild type plastomes

**Figure S1.**
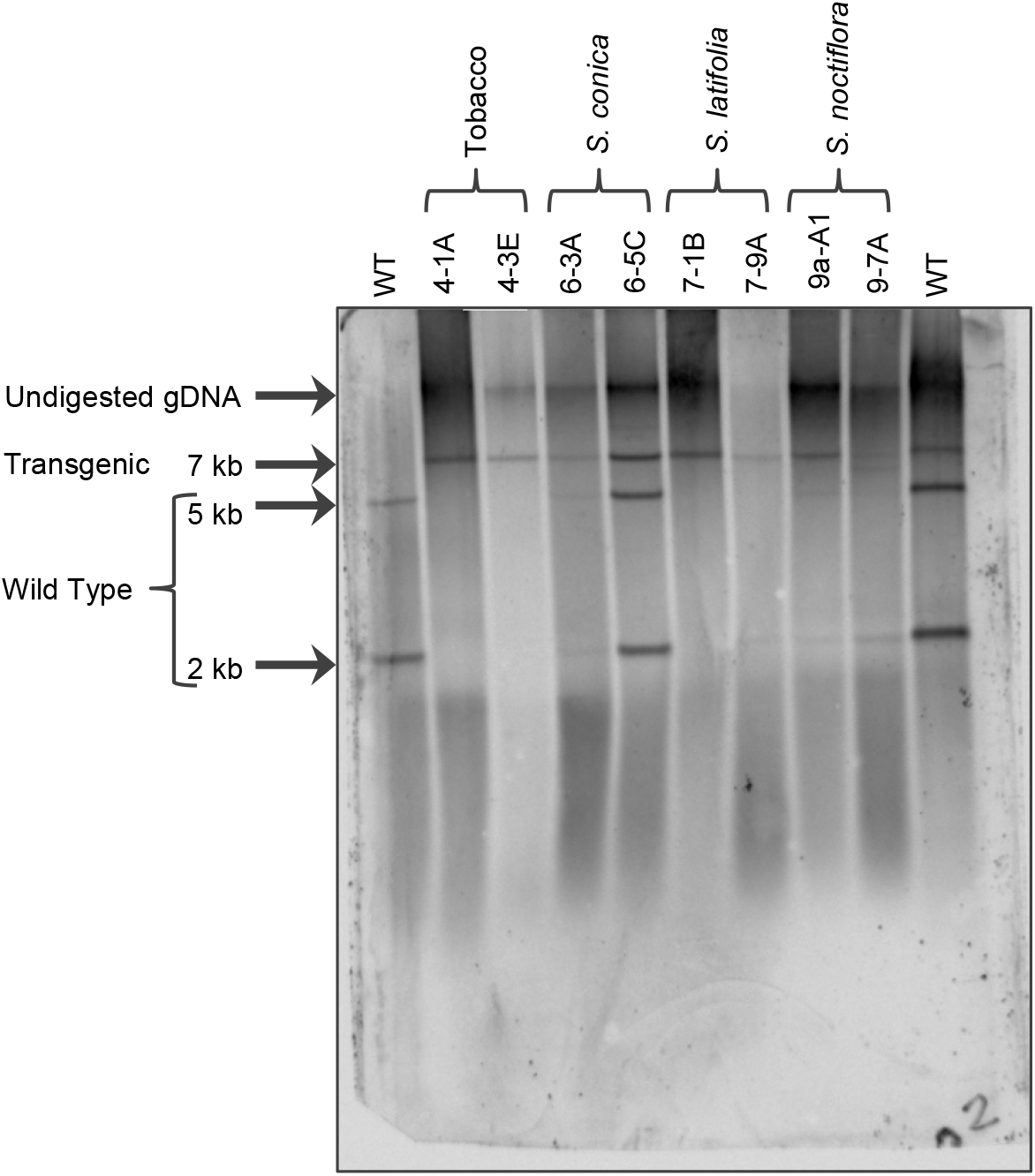
Southern blot analysis of select wild type and transgenic lines, confirming presence of the transgenic construct.

**Figure S2.**
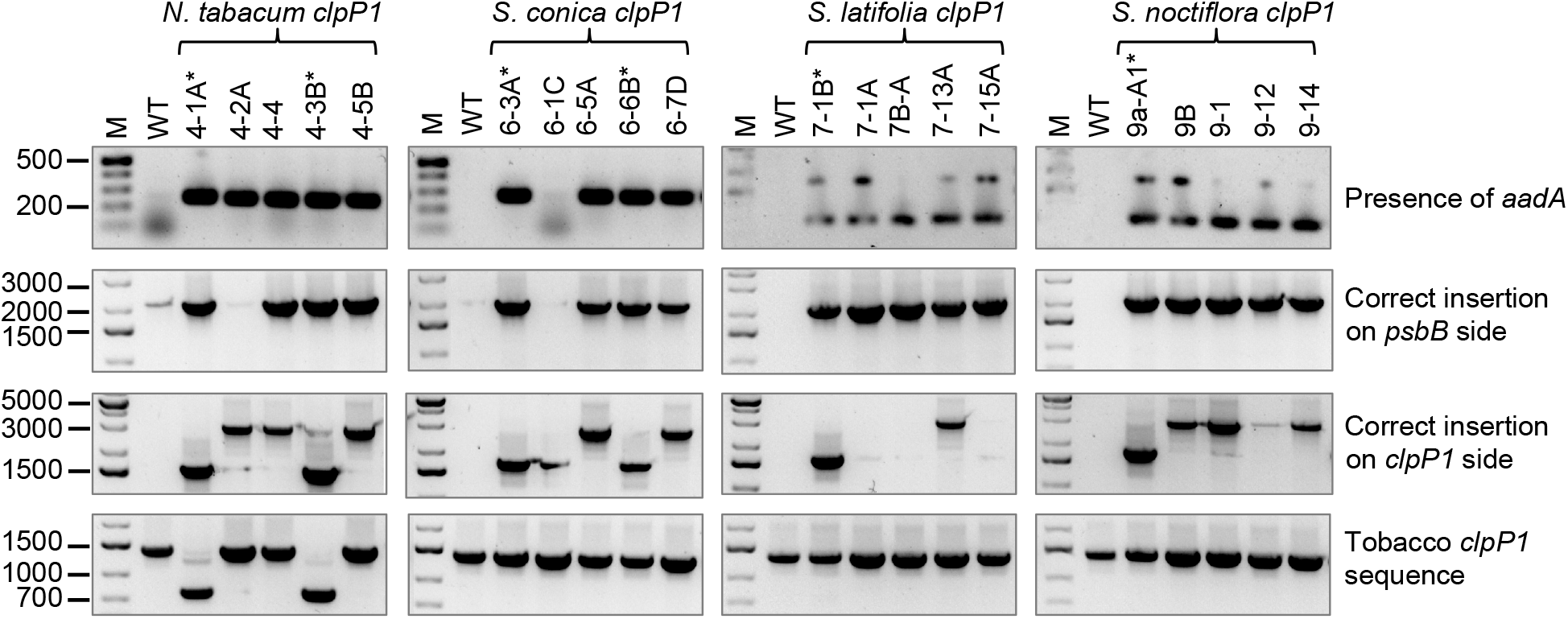
Examples of transgenic lines with improper insertions or internal recombination events. The lines marked with asterisks exhibit the expected product size for each of the four PCR markers (see Figure 3).

## Notes

### Competing Interest Statement

The authors have declared no competing interest.

## REFERENCES

Adrion JR, White PS, Montooth KL. 2016. The roles of compensatory evolution and constraint in aminoacyl tRNA synthetase evolution. Molecular Biology and Evolution 33:152.

Aguileta G, Refregier G, Yockteng R, Fournier E, Giraud T. 2009. Rapidly evolving genes in pathogens: methods for detecting positive selection and examples among fungi, bacteria, viruses and protists. Infection, Genetics and Evolution 9:656–670.

Apitz J, Nishimura K, Schmied J, Wolf A, Hedtke B, van Wijk KJ, Grimm B. 2016. Posttranslational control of ALA synthesis includes GluTR degradation by Clp protease and stabilization by GluTR-binding protein. Plant Physiology 170:2040–2051.

Barnard-Kubow KB, So N, Galloway LF. 2016. Cytonuclear incompatibility contributes to the early stages of speciation. Evolution 70:2752–2766.

Bock R. 2001. Transgenic plastids in basic research and plant biotechnology. Journal of Molecular Biology 312:425–438.

Bogdanova VS, Zaytseva OO, Mglinets AV, Shatskaya NV, Kosterin OE, Vasiliev GV. 2015. Nuclear-cytoplasmic conflict in pea (Pisum sativum L.) is associated with nuclear and plastidic candidate genes encoding acetyl-CoA carboxylase subunits. PloS one 10:e0119835.

Dittmar D, Reder A, Schlüter R, Riedel K, Hecker M, Gerth U. 2020. Complementation studies with human ClpP in Bacillus subtilis. Biochimica et Biophysica Acta (BBA)-Molecular Cell Research 1867:118744.

Doyle JJ, Doyle JL. 1990. Isolation of plant DNA from fresh tissue. Focus 12:39–40.

Erixon P, Oxelman B. 2008. Whole-gene positive selection, elevated synonymous substitution rates, duplication, and indel evolution of the chloroplast clpP1 gene. PloS one 3:e1386.

Fajardo D, Senalik D, Ames M, Zhu H, Steffan SA, Harbut R, Polashock J, Vorsa N, Gillespie E, Kron K. 2013. Complete plastid genome sequence of Vaccinium macrocarpon: structure, gene content, and rearrangements revealed by next generation sequencing. Tree Genetics & Genomes 9:489–498.

Forsythe ES, Williams AM, Sloan DB. 2021. Genome-wide signatures of plastid-nuclear coevolution point to repeated perturbations of plastid proteostasis systems across angiosperms. Plant Cell In Press.

Fujii S, Bond CS, Small ID. 2011. Selection patterns on restorer-like genes reveal a conflict between nuclear and mitochondrial genomes throughout angiosperm evolution. Proceedings of the National Academy of Sciences of the United States of America 108:1723–1728.

Gould SB, Waller RF, McFadden GI. 2008. Plastid evolution. Annual Review of Plant Biology 59:491–517.

Gray MW. 2012. Mitochondrial evolution. Cold Spring Harbor Perspectives in Biology 4:a011403.

Greiner S, Golczyk H, Malinova I, Pellizzer T, Bock R, Börner T, Herrmann RG. 2020. Chloroplast nucleoids are highly dynamic in ploidy, number, and structure during angiosperm leaf development. The Plant Journal 102:730–746.

Greiner S, Rauwolf U, Meurer J, Herrmann RG. 2011. The role of plastids in plant speciation. Molecular ecology 20:671–691.

Haberle RC, Fourcade HM, Boore JL, Jansen RK. 2008. Extensive rearrangements in the chloroplast genome of Trachelium caeruleum are associated with repeats and tRNA genes. Journal of Molecular Evolution 66:350–361.

Hajdukiewicz PT, Allison LA, Maliga P. 1997. The two RNA polymerases encoded by the nuclear and the plastid compartments transcribe distinct groups of genes in tobacco plastids. EMBO Journal 16:4041–4048.

Havird JC, Forsythe ES, Williams AM, Werren JH, Dowling DK, Sloan DB. 2019. Selfish mitonuclear conflict. Current Biology 29:R496–R511.

Havird JC, Trapp P, Miller C, Bazos I, Sloan DB. 2017. Causes and consequences of rapidly evolving mtDNA in a plant lineage. Genome Biology and Evolution 9:323–336.

Hirao T, Watanabe A, Kurita M, Kondo T, Takata K. 2008. Complete nucleotide sequence of the Cryptomeria japonica D. Don. chloroplast genome and comparative chloroplast genomics: diversified genomic structure of coniferous species. BMC Plant Biology 8:70.

Huang C, Wang S, Chen L, Lemieux C, Otis C, Turmel M, Liu X-Q. 1994. The Chlamydomonas chloroplast clpP gene contains translated large insertion sequences and is essential for cell growth. Molecular and General Genetics 244:151–159.

Hughes AL, Nei M. 1988. Pattern of nucleotide substitution at major histocompatibility complex class I loci reveals overdominant selection. Nature 335:167–170.

Jafari F, Zarre S, Gholipour A, Eggens F, Rabeler RK, Oxelman B. 2020. A new taxonomic backbone for the infrageneric classification of the species-rich genus Silene (Caryophyllaceae). Taxon 69:337–368.

Kanevski I, Maliga P, Rhoades DF, Gutteridge S. 1999. Plastome engineering of ribulose-1, 5-bisphosphate carboxylase/oxygenase in tobacco to form a sunflower large subunit and tobacco small subunit hybrid. Plant Physiology 119:133–142.

Kim J, Olinares PD, Oh S-h, Ghisaura S, Poliakov A, Ponnala L, van Wijk KJ. 2013. Modified Clp protease complex in the ClpP3 null mutant and consequences for chloroplast development and function in Arabidopsis. Plant Physiology 162:157–179.

Kim J, Rudella A, Rodriguez VR, Zybailov B, Olinares PDB, van Wijk KJ. 2009. Subunits of the plastid ClpPR protease complex have differential contributions to embryogenesis, plastid biogenesis, and plant development in Arabidopsis. Plant Cell 21:1669–1692.

Koussevitzky S, Stanne TM, Peto CA, Giap T, Sjögren LL, Zhao Y, Clarke AK, Chory J. 2007. An Arabidopsis thaliana virescent mutant reveals a role for ClpR1 in plastid development. Plant Molecular Biology 63:85–96.

Kuroda H, Maliga P. 2003. The plastid clpP1 protease gene is essential for plant development. Nature 425:86–89.

Majeran W, Wollman F-A, Vallon O. 2000. Evidence for a role of ClpP in the degradation of the chloroplast cytochrome b6f complex. Plant Cell 12:137–149.

Maliga P, Moll B, Svab Z. 1990. Toward manipulation of plastid genes in higher plants. In: Zelitch I, editor. Perspectives in genetic and biochemical regulation of photosynthesis. New York. p. 133–143.

Montandon C, Friso G, Liao J-YR, Choi J, van Wijk KJ. 2019. In Vivo Trapping of Proteins Interacting with the Chloroplast CLPC1 Chaperone: Potential Substrates and Adaptors. Journal of proteome research 18:2585–2600.

Moreno JC, Martínez-Jaime S, Schwartzmann J, Karcher D, Tillich M, Graf A, Bock R. 2018. Temporal proteomics of inducible RNAi lines of Clp protease subunits identifies putative protease substrates. Plant Physiology 176:1485–1508.

Moreno JC, Tiller N, Diez M, Karcher D, Tillich M, Schöttler MA, Bock R. 2017. Generation and characterization of a collection of knock-down lines for the chloroplast Clp protease complex in tobacco. Journal of Experimental Botany 68:2199–2218.

Nishimura K, Apitz J, Friso G, Kim J, Ponnala L, Grimm B, van Wijk KJ. 2015. Discovery of a unique Clp component, ClpF, in chloroplasts: a proposed binary ClpF-ClpS1 adaptor complex functions in substrate recognition and delivery. Plant Cell 27:2677–2691.

Nishimura K, Asakura Y, Friso G, Kim J, Oh S-h, Rutschow H, Ponnala L, van Wijk KJ. 2013. ClpS1 is a conserved substrate selector for the chloroplast Clp protease system in Arabidopsis. Plant Cell 25:2276–2301.

Nishimura K, van Wijk KJ. 2015. Organization, function and substrates of the essential Clp protease system in plastids. Biochimica et Biophysica Acta-Bioenergetics 1847:915–930.

Nováková E, Zablatzká L, Brus J, Nesrstová V, Hanáček P, Kalendar R, Cvrčková F, Majeský Ĺ, Smýkal P. 2019. Allelic diversity of acetyl coenzyme A carboxylase accD/bccp genes implicated in nuclear-cytoplasmic conflict in the wild and domesticated pea (Pisum sp.). International Journal of Molecular Sciences 20:1773.

Osada N, Akashi H. 2012. Mitochondrial-nuclear interactions and accelerated compensatory evolution: evidence from the primate cytochrome C oxidase complex. Molecular Biology and Evolution 29:337.

Pett W, Lavrov DV. 2015. Cytonuclear interactions in the evolution of animal mitochondrial tRNA metabolism. Genome Biology and Evolution 7:2089–2101.

Pulido P, Llamas E, Llorente B, Ventura S, Wright LP, Rodríguez-Concepción M. 2016. Specific Hsp100 chaperones determine the fate of the first enzyme of the plastidial isoprenoid pathway for either refolding or degradation by the stromal Clp protease in Arabidopsis. PLoS Genetics 12:e1005824.

Rand DM, Haney RA, Fry AJ. 2004. Cytonuclear coevolution: the genomics of cooperation. Trends in Ecology & Evolution 19:645–653.

Rautenberg A, Sloan DB, Aldén V, Oxelman B. 2012. Phylogenetic relationships of Silene multinervia and Silene section Conoimorpha (Caryophyllaceae). Systematic Botany 37:226–237.

Rockenbach KD, Havird JC, Monroe JG, Triant DA, Taylor DR, Sloan DB. 2016. Positive selection in rapidly evolving plastid-nuclear enzyme complexes. Genetics 204:1507–1522.

Roger AJ, Muñoz-Gómez SA, Kamikawa R. 2017. The origin and diversification of mitochondria. Current Biology 27:R1177–R1192.

Rousseau-Gueutin M, Ayliffe MA, Timmis JN. 2011. Conservation of plastid sequences in the plant nuclear genome for millions of years facilitates endosymbiotic evolution. Plant Physiology 157:2181–2193.

Schmitz-Linneweber C, Kushnir S, Babiychuk E, Poltnigg P, Herrmann RG, Maier RM. 2005. Pigment deficiency in nightshade/tobacco cybrids is caused by the failure to edit the plastid ATPase alpha-subunit mRNA. Plant Cell 17:1815–1828.

Shikanai T, Shimizu K, Ueda K, Nishimura Y, Kuroiwa T, Hashimoto T. 2001. The chloroplast clpP gene, encoding a proteolytic subunit of ATP-dependent protease, is indispensable for chloroplast development in tobacco. Plant & Cell Physiology 42:264–273.

Sjögren LL, Stanne TM, Zheng B, Sutinen S, Clarke AK. 2006. Structural and functional insights into the chloroplast ATP-dependent Clp protease in Arabidopsis. Plant Cell 18:2635–2649.

Sloan DB, Havird JC, Sharbrough J. 2017. The on-again, off-again relationship between mitochondrial genomes and species boundaries. Molecular ecology 26:2212–2236.

Sloan DB, Triant DA, Forrester NJ, Bergner LM, Wu M, Taylor DR. 2014. A recurring syndrome of accelerated plastid genome evolution in the angiosperm tribe Sileneae (Caryophyllaceae). Molecular phylogenetics and evolution 72:82–89.

Sloan DB, Triant DA, Wu M, Taylor DR. 2014. Cytonuclear interactions and relaxed selection accelerate sequence evolution in organelle ribosomes. Molecular Biology and Evolution 31:673–682.

Sloan DB, Warren JM, Williams AM, Wu Z, Abdel-Ghany SE, Chicco AJ, Havird JC. 2018. Cytonuclear integration and co-evolution. Nature Reviews Genetics 19:635–648.

Sobanski J, Giavalisco P, Fischer A, Kreiner JM, Walther D, Schöttler MA, Pellizzer T, Golczyk H, Obata T, Bock R. 2019. Chloroplast competition is controlled by lipid biosynthesis in evening primroses. Proceedings of the National Academy of Sciences 116:5665–5674.

Straub SC, Fishbein M, Livshultz T, Foster Z, Parks M, Weitemier K, Cronn RC, Liston A. 2011. Building a model: developing genomic resources for common milkweed (Asclepias syriaca) with low coverage genome sequencing. BMC Genomics 12:211.

Stupar RM, Lilly JW, Town CD, Cheng Z, Kaul S, Buell CR, Jiang J. 2001. Complex mtDNA constitutes an approximate 620-kb insertion on Arabidopsis thaliana chromosome 2: implication of potential sequencing errors caused by large-unit repeats. Proceedings of the National Academy of Sciences of the United States of America 98:5099–5103.

Svab Z, Hajdukiewicz P, Maliga P. 1990. Stable transformation of plastids in higher plants. Proceedings of the National Academy of Sciences 87:8526–8530.

Svab Z, Maliga P. 1993. High-frequency plastid transformation in tobacco by selection for a chimeric aadA gene. Proceedings of the National Academy of Sciences 90:913–917.

Tapken W, Kim J, Nishimura K, van Wijk KJ, Pilon M. 2015. The Clp protease system is required for copper ion-dependent turnover of the PAA 2/HMA 8 copper transporter in chloroplasts. New Phytologist 205:511–517.

Touzet P, Budar F. 2004. Unveiling the molecular arms race between two conflicting genomes in cytoplasmic male sterility? Trends in plant science 9:568–570.

Welsch R, Zhou X, Yuan H, Álvarez D, Sun T, Schlossarek D, Yang Y, Shen G, Zhang H, Rodriguez-Concepcion M. 2018. Clp protease and OR directly control the proteostasis of phytoene synthase, the crucial enzyme for carotenoid biosynthesis in Arabidopsis. Molecular Plant 11:149–162.

Weng ML, Ruhlman TA, Jansen RK. 2016. Plastid-nuclear interaction and accelerated coevolution in plastid ribosomal genes in Geraniaceae. Genome Biology and Evolution 8:1824–1838.

Williams AM, Friso G, van Wijk KJ, Sloan DB. 2019. Extreme variation in rates of evolution in the plastid Clp protease complex. Plant Journal 98:243–259.

Williams AV, Boykin LM, Howell KA, Nevill PG, Small I. 2015. The complete sequence of the Acacia ligulata chloroplast genome reveals a highly divergent clpP1 gene. PloS one 10:e0125768.

Wu G-Z, Chalvin C, Hoelscher M, Meyer EH, Wu XN, Bock R. 2018. Control of retrograde signaling by rapid turnover of GENOMES UNCOUPLED1. Plant Physiology 176:2472–2495.

Yan Z, Ye G, Werren J. 2019. Evolutionary rate correlation between mitochondrial-encoded and mitochondria-associated nuclear-encoded proteins in insects. Molecular Biology and Evolution 36:1022–1036.

Yang Z. 2007. PAML 4: Phylogenetic Analysis by Maximum Likelihood. Molecular Biology and Evolution 24:1586–1591.

Yao X, Tang P, Li Z, Li D, Liu Y, Huang H. 2015. The first complete chloroplast genome sequences in Actinidiaceae: genome structure and comparative analysis. PloS one 10:e0129347.

Zhang J, Ruhlman TA, Sabir J, Blazier JC, Jansen RK. 2015. Coordinated rates of evolution between interacting plastid and nuclear genes in Geraniaceae. The Plant Cell 27:563–573.

Zhang J, Ruhlman TA, Sabir JS, Blazier JC, Weng ML, Park S, Jansen RK. 2016. Coevolution between nuclear-encoded DNA replication, recombination, and repair genes and plastid genome complexity. Genome Biology and Evolution 8:622–634.

Zhang Y, Ma J, Yang B, Li R, Zhu W, Sun L, Tian J, Zhang L. 2014. The complete chloroplast genome sequence of Taxus chinensis var. mairei (Taxaceae): loss of an inverted repeat region and comparative analysis with related species. Gene 540:201–209.

Zheng B, MacDonald TM, Sutinen S, Hurry V, Clarke AK. 2006. A nuclear-encoded ClpP subunit of the chloroplast ATP-dependent Clp protease is essential for early development in Arabidopsis thaliana. Planta 224:1103–1115.

Zupok A, Kozul D, Schöttler MA, Niehörster J, Garbsch F, Liere K, Malinova I, Bock R, Greiner S. 2020. A photosynthesis operon in the chloroplast genome drives speciation in evening primroses. bioRxiv:2020.2007.2003.186627.

